# Encephalomyocarditis virus non-structural protein 2C degrades NDP52 autophagy protein to promote its own survival

**DOI:** 10.1101/2024.12.13.628388

**Authors:** Rongqian Mo, Rongrong Cheng, Pingan Dong, Tingting Ma, Yaxin Zhang, Jingying Xie, Shasha Li, Huixia Li, Adi Idris, Xiangrong Li, Ruofei Feng

## Abstract

Encephalomyocarditis virus (EMCV) is an important zoonotic pathogen that causes encephalitis and myocarditis, primarily in pigs and in some mammals. Nuclear dot protein (NDP) 52 is an important autophagy adaptor protein that is known to target microbial pathogens, including viruses into autophagosomes to facilitate the selective autophagy process. Here, we wanted to investigate how an important zoonotic virus such as EMCV interacts with NDP52. We found that NDP52 negatively regulates the entry and replication phases of EMCV and interacts with EMCV VP1/VP2 proteins to mediate its autophagic degradation. EMCV counteracts this by exerting autophagy induction through its encoded 2C protein. The autophagy machinery then hijacks NDP52 and transports it to lysosomes for subsequent degradation via late endosomal molecules Rab7 and Rab9. Our study describes a novel mechanism by which EMCV escapes the host’s antiviral response to promote its survival by hijacking the autophagy pathway using the non-structural protein 2C of EMCV.

**Importance:** EMCV is an important zoonotic pathogen that can induce autophagy, but the regulatory role of autophagy receptors in EMCV infection has been rarely reported. Herein, we show that NDP52 is a novel host limiting factor for EMCV, which not only inhibits early EMCV endocytosis, but also restricts EMCV replication in vitro by controlling autophagic degradation of EMCV structural proteins VP1 and VP2. In order to counteract this, EMCV uses the N-terminal region of its non-structural protein 2C to interact with NDP52, and hijacks the autophagy machine to transport NDP52 to lysosomes for degradation, which weakens NDP52-mediated degradation of VP1 and VP2. Importantly, we reveal a novel mechanism by which EMCV evades the host antiviral response, contributing to a deeper understanding of the infectious mechanism of EMCV and providing new ideas for the design of antiviral drug targets.

## Introduction

Encephalomyocarditis virus (EMCV) causes severe encephalitis and myocarditis in animals (1). In the veterinary world, EMCV is an important cause of acute myocarditis in piglets and of fetal death or abortion in pregnant sows. EMCV belongs to the *Cardiomyovirus* genus and is a single-stranded positive-stranded non-enveloped RNA virus. Its 7.8 kb genome encodes four structural proteins (VP1, VP2, VP3 and VP4), seven non-structural proteins (2A^pro^, 2B, 2C, 3A, 3B, 3C^pro^ and 3D) and a virally encoded precursor protein, L (2). The structural proteins are mainly involved in viral entry and assembly of viral particles and are home to viral antigenic sites. Among them, VP1 is the most antigenic, and is an important protein for the topology, antigenicity, receptor adsorption, and decapsulation of the surface of virus particles (2). We were the first to show that EMCV VP1 and VP2 inhibit the type I interferon (IFN) antiviral responses to promote EMCV proliferation (3). Non-structural proteins mainly encode viral genome replication and protein processing related proteins, which are associated with viral replication, infection and pathogenesis (4). Among these non-structural proteins, 2C possesses ATPase activity and a deconjugating enzyme structural domain. 2C can associate with 2B non-structural protein to rearrange host cell membrane and vesicle formation through the regulation of Ca^2+^ content in the endoplasmic reticulum (5-7). 2C can also bind to the 3’ untranslated region (UTR) of the EMCV genome to participate in viral RNA replication. Importantly, EMCV 2C protein can induce cellular autophagy (8).

Autophagy is an evolutionarily cellular conserved process of degrading its own cytoplasmic proteins and damaged organelles using lysosomes under the regulation of autophagy-related genes (ATGs) through the formation of autophagosomes and autophagic lysosomes (9). As opposed to non-selective autophagic process of protein degradation for the purpose of maintaining basic material energy metabolism, autophagy receptors have been shown to specifically recognize proteins, including those encoded by pathogens, to autophagosomes for degradation (selective autophagy) (10). Hence, this selective autophagic process can functionally dampen viral propagation and dissemination in the host. NDP52 also known as calcium binding and coiled-coil domain 2 (CALCOCO2), is a known CALCOCO family of proteins associated with selective autophagy (11). NDP52 has been shown to recognize and transport ubiquitinated porcine epidemic diarrhea virus (PEDV) nucleocapsid N protein for eventual autolysosomal degradation (12-14). Another autophagic receptor, sequestosome 1/polyubiquitin-binding protein p62 (SQSTM1/p62), has been shown to recognize proteins from a range of viruses including, Seneca Valley virus (SVV) (15), dengue virus (DENV) (16) and Zika virus (ZIKV) (17), for its eventual degradation to result in the dampening of viral proliferation. Armed with this knowledge, we speculate that this viral interplay with the host autophagic machinery exists within the context of an EMCV infection. Hence, this detailed study seeks to elucidate this undefined mechanism.

In this study, we reveal an antiviral defense function of NDP52 in response to EMCV infection. We demonstrate that EMCV 2C protein is an autophagy blocker that directly induces the degradation of NDP52 through the Rab7/Rab9-dependent autophagic lysosomal pathway. This alleviates the antiviral function of NDP52 to allow an environment conducive for EMCV infection to thrive.

## Results

### EMCV infection down-regulates NDP52 protein expression

EMCV infection has been shown to initiate the onset of autophagy (8). However, the mechanistic details of this are not well characterized. Consistent with previous findings (8), EMCV infection promoted the expression of the autophagy-regulated gene ATG7, the conversion of the unlipidated light chain 3 (LC3) B-I to the phosphatidylethanolamine lipidated LC3B-II form, a hallmark of autophagy, and the decrease of the expression of NDP52 and SQSTM1/p62 (Fig. 1A). Importantly, NDP52 expression decreases over time post-EMCV infection at the protein (Fig. 1B and D), not transcriptional level (Fig. 1C and E), irrespective of the infectious dose. Based on this we postulated that EMCV is directly degrading NDP52. Indeed, immunofluorescence analysis reveals a direct co-localization between NDP52 and EMCV (VP1) (Fig. 1F). Correlating with the decreased NDP52 fluorescence signal in EMCV infected cells (Fig. 1F), we also observed an EMCV infectious dose-dependent decrease in NDP52 levels (Fig. 1G). This further strengthens the direct link between EMCV infectivity and degradation of the NDP52 autophagic adaptor protein.

**Fig 1.**
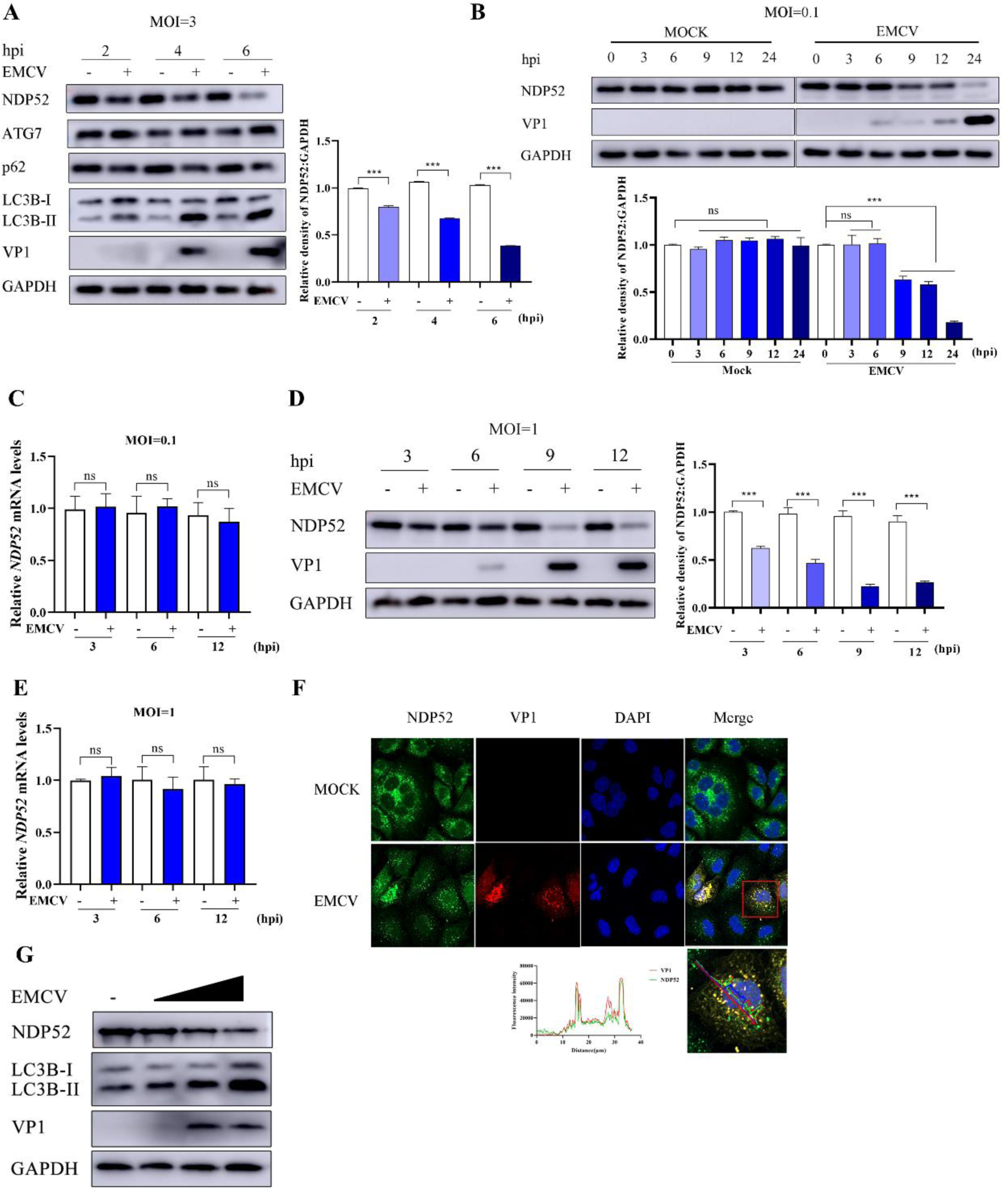
EMCV infection down-regulates NDP52 protein expression. (A, B, D) A549 cells were infected or uninfected with EMCV for 2, 4, 6 h/0, 3, 6, 9, 12, 24 h/3, 6, 9, 12 h at MOI of 3/0.1/1. Western blotting was performed to examine the expression of NDP52 using an anti-NDP52 antibody, EMCV capsid protein VP1. GAPDH served as a loading control. The gray ratios of NDP52/GAPDH in Figure (A, B, D) were analyzed using Image J software, respectively. Data are accumulated from three independent experiments. (C, E) The mRNA expression of NDP52 following EMCV infection. A549 cells were infected or uninfected with EMCV for 0, 6, and 12 h (MOI = 0.1/1). Real-time quantitative RT-PCR was used to analyze the transcriptional level of NDP52 and normalized to GAPDH mRNA. Statistical data from three independent infection experiments is shown. (F) A549 cells were infected or uninfected with EMCV for 4 h (MOI=3). The cells were stained with NDP52 antibodies, VP1 antibodies, and 4’,6-diamidino-2-phenylindole (DAPI), then examined by confocal microscopy. Samples were stained with NDP52 antibody (green), VP1 antibody (red) and 4’,6-diamidino-2-phenylindole (DAPI) (blue), then examined by confocal microscopy. Scale bar: 20 μm. (G) HEK-293 cells were infected or uninfected with EMCV (MOI=0.1, 1, 2) for 12 h. Western blotting was performed to examine the expression of NDP52 using an anti-NDP52 antibody, EMCV capsid protein VP1. GAPDH served as a loading control. The data were represented as the mean ± SD of three independent experiments. The measurements were performed in technical duplicate. Statistical significance was denoted as \*\**P* < 0.01, and \*\*\**P* < 0.001.

### EMCV degrades NDP52 via the Rab7/Rab9-dependent autophagic lysosomal pathway

There are several host cellular pathways that eventuate in target protein degradation, including the ubiquitin-proteasome, autophagy lysosomal and apoptotic pathways. We want to explore which of these pathways lead to EMCV infection-driven NDP52 degradation. Pretreatment of cells with either proteasome (MG132), autophagy (chloroquine, CQ), or pan-caspase apoptosis (Z-VAD-FMK) pathway inhibitors before EMCV infection revealed that only CQ could revert EMCV-mediated degradation of NDP52 (Fig. 2A) and in a dose-dependent manner (Fig. 2B). Treatment of EMCV infected cells with early and late autophagy inhibitors, 3-MA and Baf.A1, and the lysosomal inhibitor, NH_4_Cl, just Baf.A1and NH_4_Cl revived NDP52 levels (Fig. 2C and D). Treatment with an intracellular protein synthesis inhibitor, cyclohexamide (CHX), failed to rescue EMCV infection-mediated NDP52 degradation (Figure 2E), further confirming that the degradation of NDP52 is occurring via the autophagic lysosomal pathway post-EMCV infection. The enhanced colocalization of NDP52 with lysosomal structures in EMCV infected cells (Fig. 2F) indicates the involvement of the lysosomal pathway. Late endosomal Rab7/Rab9 proteins are known to target biomolecules, such as proteins for lysosomal catabolism (18-21). Endosomal function has been shown to intertwine with the cellular autophagic machinery. Importantly, Rab9 can form a complex with NDP52, which mediates the movement of NDP52 and viral proteins to the lysosome for degradation (22). Indeed, when Rab7 and 9 were knocked down EMCV-mediated NDP52 degradation was reversed (Fig. 2G). In summary, we show that EMCV degrades NDP52 via the Rab7/Rab9-dependent autophagic lysosomal pathway.

**Fig 2.**
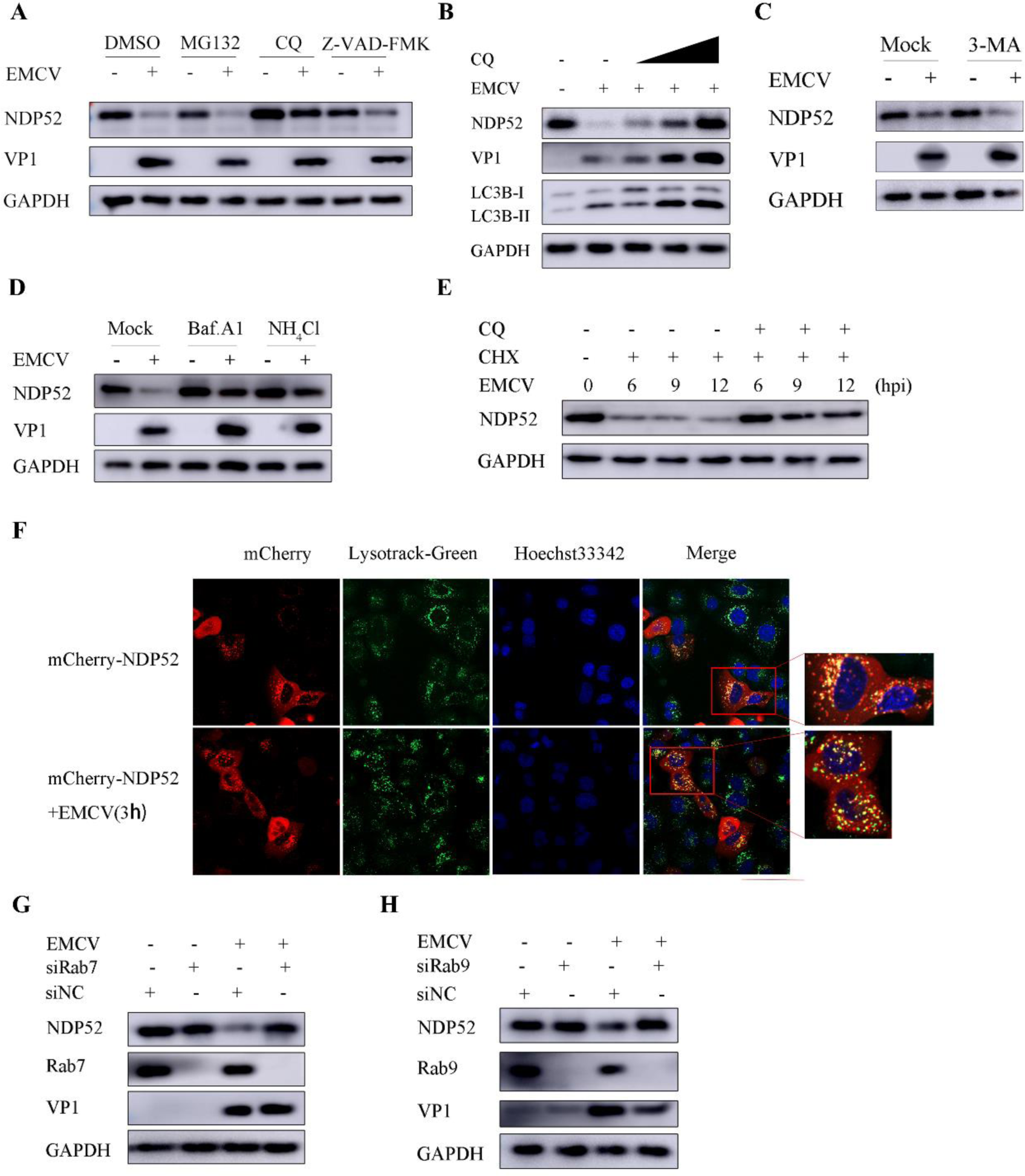
EMCV degrades NDP52 via the Rab7/Rab9-dependent autophagic lysosomal pathway. (A)A549 cells were infected or uninfected with EMCV for 2 h (MOI=1) and then treated for 7 h with dimethyl sulfoxide (DMSO), MG132 (10 µM), CQ (50 µM), or Z-VAD-FMK (20 µM), respectively. Western blotting was used to detect the expression of NDP52 protein and VP1. GAPDH was used as a loading control. (B) A549 cells were infected or uninfected with EMCV for 2 h (MOI=1) and then treated for 7 h with CQ (25 µM, 50 μM, 100 μM), respectively, and western blotting was used to detect the expression of NDP52 protein and VP1. GAPDH was used as a loading control. (C, D) A549 cells were infected or uninfected with EMCV for 2 h (MOI=1) and then treated for 7 h with PBS, 3-MA (20 mM), Bafilomycin A1 (1 mM), or NH_4_Cl (20 mM), respectively. Western blotting was used to detect the expression of NDP52 protein and VP1. GAPDH was used as a loading control. (E) A549 cells were infected or uninfected with EMCV for 2 h (MOI=1), then treated for 0, 6, 9, 12 h with cycloheximide (CHX) (100 µg/ml), dimethyl sulfoxide (DMSO), CQ (50 µM), respectively. Western blotting was used to detect the expression of NDP52 protein and VP1. GAPDH was used as a loading control. (F)A549 cells were transfected with 1.0 µg of mCherrry-NDP52 plasmid for 24 h, then infected or uninfected with EMCV for 3 h (MOI=3). The cells were stained with Lyso-Tracker Green for 1h and Hoechst 33342 for 10min, followed by fluorescence analysis. Scale bar = 10 μm. (G, H, I) A549 cells were transfected with 20mM of siNC or siRNA (siRab7, siRab9) for 36 h. And then infected or uninfected with EMCV for 9 h (MOI=1). Western blotting was used to detect the expression of NDP52 protein, Rab7/rab9 and VP1. GAPDH was used as a loading control.

### EMCV 2C and 3C are key viral proteins involved in NDP52 degradation

We next wanted to determine which EMCV proteins are directly responsible for NDP52 degradation. Screening of several EMCV structural and non-structural proteins revealed that only 2C and 3C overexpression was able to result in NDP52 degradation (Fig. 3A). Indeed, genetic silencing of 2C or 3C with siRNAs reversed EMCV-driven NDP52 degradation (Fig. 3B-G). More importantly, co-immunoprecipitation (co-IP) (Fig. 3H) and confocal microscopic analyses (Fig. 3J) showed that EMCV 2C and 3C can directly interact with NDP52. This is consistent with previous work that showed that CVB3 3C can functionally cleave NDP52 through its protease activity to dampen autophagic function (23-25). From here forward, we focused our attention on the less studied EMCV 2C protein to mechanistically uncover how it degrades NDP52.

**Fig 3.**
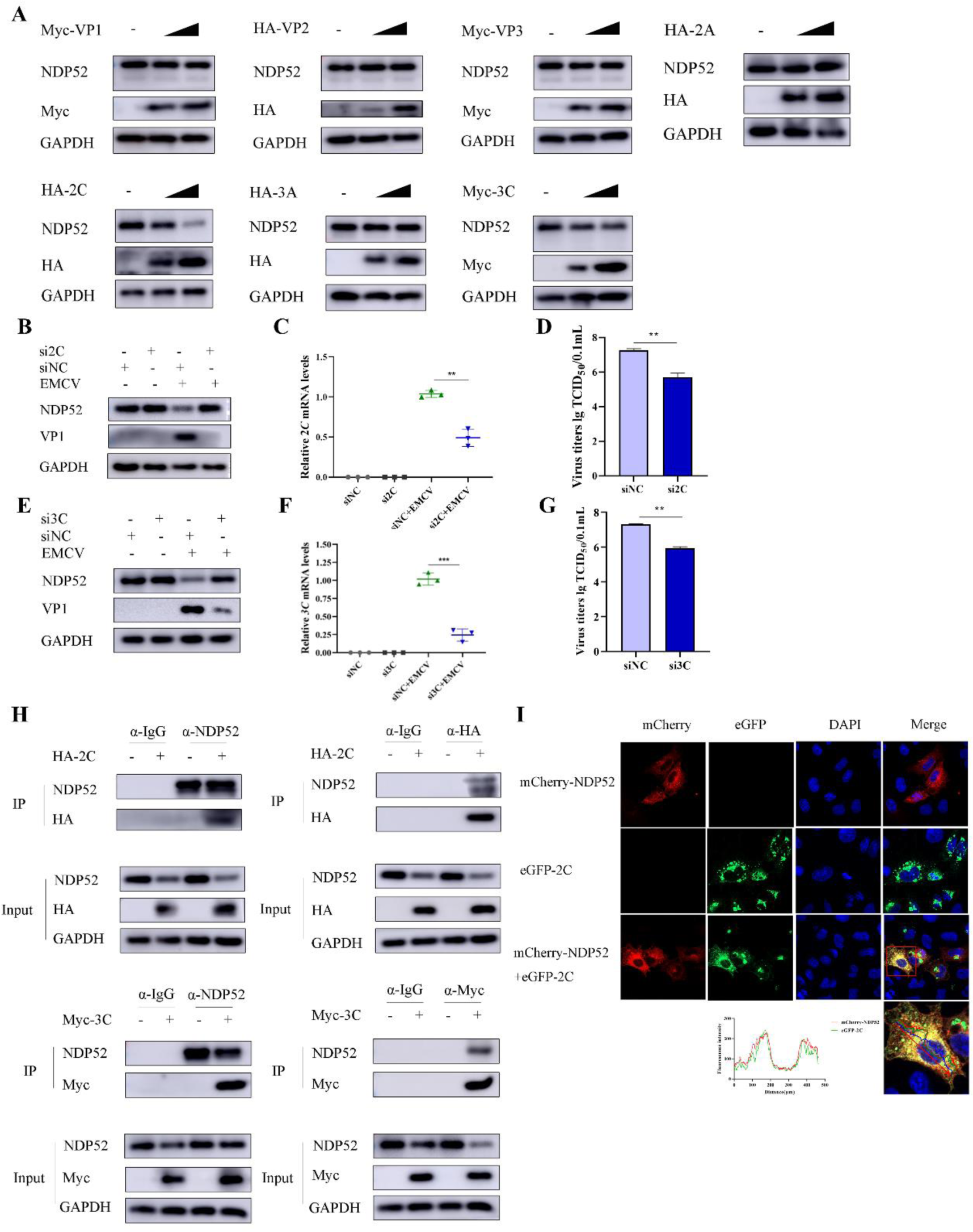
EMCV 2C and 3C are key viral proteins involved in NDP52 degradation. (A)A549 cells were transfected with 1.0 µg of EV or Myc-VP1, HA-VP2, Myc-VP3, HA-VP2, HA-2A, HA-2C, HA-3A, Myc-3C plasmids for 36 h, then the cells were collected using western blotting to detect expression NDP52, Myc-tagged VP1/VP3/3C proteins, HA-tagged VP2/2A/2C/3A proteins, GAPDH was used as a loading control. (B, E)A549 cells were transfected with 20 mM of siNC or siRNA (si2C,si3C) for 36 h followed by EMCV (MOI = 1) infect for 9 h. Western blotting was used to detect the protein expression of NDP52, VP1, GAPDH was used as a loading control. (C, F) A549 cells were transfected with 20mM of siNC or siRNA (si2C,si3C) for 36 h followed by EMCV (MOI = 1) infection for 9 h. Total cellular RNA was extracted, and then the mRNA transcript levels of 2C and 3C were detected by qRT-PCR, and normalized to GAPDH mRNA. Statistical data from three independent infection experiments is shown. (*, *P* < 0.05; ***, *P* < 0.001; NS, not significant). (D, G) A549 cells were transfected with 20mM of siNC or siRNA (si2C,si3C) for 36 h followed by EMCV (MOI = 1) infection for 9 h. Samples were subjected to TCID_50_ assay.(*, *P* < 0.05; ***, *P* < 0.001; NS, not significant). (H, J) A549 cells were transfected with 1.0 µg of EV or HA-2C/Myc-3C plasmids for 36h.Cell samples were prepared for Co-IP analysis. (I) A549 cells were cotransfected with 1.0μg mCherry-NDP52, eGFP-2C plasmids for 24 h. Samples were stained with DAPI (blue), then examined by confocal microscopy. Scale bar: 20 μm.

### EMCV 2C degrades NDP52 via the autophagic lysosomal pathway

We further confirmed the role of EMCV 2C in NDP52 autophagic lysosomal degradation by exogenously overexpressing EMCV 2C in cells. Consistent with our preliminary findings (Fig. 3), EMCV 2C was able to degrade both endogenous and exogenous NDP52 and that this effect can be reversed using the autophagy inhibitor, CQ (Fig. 4A and B). Moreover, increasing amounts of exogenously added 2C can promote autophagy by virtue of intracellular LC3B-I to LC3B-II conversion (Fig. 4C). Importantly, NDP52 degradation was specific to EMCV 2C protein, not 2C proteins derived from other RNA viruses enterovirus 71 (EV-71), food and mouth disease virus (FMDV) and Seneca Valley virus (SVA) (Fig. 4D). Overall, we conclusively demonstrate that the degradation of NDP52 via the autophagic lysosomal pathway is seen with only exclusively with 2C protein from EMCV.

**Fig 4.**
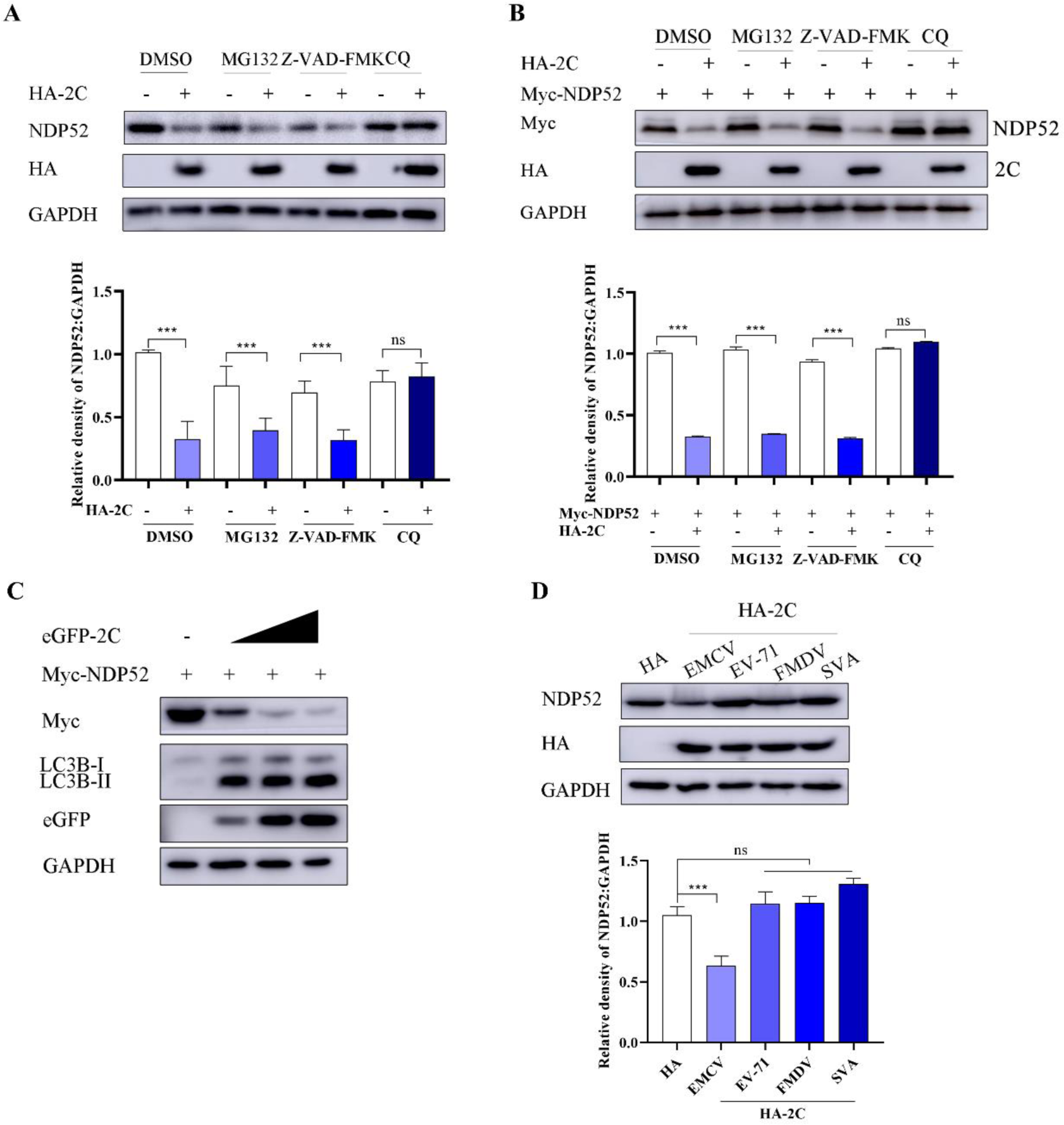
EMCV 2C degrades NDP52 via the autophagic lysosomal pathway. (A) A549 cells were transfected with 1.0 µg plasmid of HA or HA-2C for 24 h and then treated for 8 h with DMSO, MG132 (10 µM), CQ (50 µM), or Z-VAD-FMK (20 µM), respectively. Western blotting was used to detect the expression of HA-tagged 2C protein. GAPDH was used as a loading control. The gray ratios of NDP52/GAPDH in Figure A were analyzed using Image J software, respectively. (B) A549 cells were co-transfected with 1.0 µg plasmid of EV and Myc-NDP52 or co-transfected with HA-2C and Myc-NDP52 plasmids for 24 h, and then treated for 8 h with DMSO, MG132 (10 µM), CQ (50 µM), or Z-VAD-FMK (20 µM), respectively. Western blotting was used to detect the expression of NDP52, HA-tagged 2C protein. GAPDH was used as a loading control. The gray ratios of NDP52/GAPDH in Figure B were analyzed using Image J software, respectively. (C) A549 cells were transfected with 1.0 µg of EV or co-transfected with eGFP-2C (0.5µg, 1.0µg, 2.0µg) and Myc-NDP52 (1.0µg) plasmid for 36 h. Western blotting was used to detect the expression of Myc-tagged NDP52, LC3B, eGFP-tagged-2C proteins. GAPDH was used as a loading control. (D) A549 cells were transfected HA-tagged 2C of EMCV, EV-71, FMDV, and SVA. Cell lysates were collected at 36 h post-transfection (hpt), and analyzed by immunoblotting using anti-NDP52 and anti-HA antibodies. GAPDH served as a loading control. The gray ratios of NDP52/GAPDH in Figure D were analyzed using Image J software, respectively.

### EMCV 2C interacts with and degrades NDP52 via the N-terminal region

Next, we wanted to determine the biochemical interaction between EMCV 2C and NDP52 in detail. EMCV 2C is composed of an N-terminal membrane-bound structural, an intermediate ATPase domain, a zinc-finger structural domain, and a C-terminal amphipathic helical structural domain (26-28). We then generated plasmids bearing three truncated forms of 2C and found that the N-terminal region is indispensable for both endogenous (Fig. 5A) and exogenous (Fig. 5B) NDP52 degradation and that this occurs in a dose-dependent manner (Fig. 5C). Co-IP (Fig. 5D and E) and immunofluorescence co-localization analysis (Fig. 5F) also confirm that only the N-terminal region of 2C can directly interact with NDP52. Importantly, we show that the N-terminal is critical for EMCV 2C-mediated p62 degradation and LC3B-I to LC3B-II conversion (Fig. 5H). Overall, this data suggests that the N-terminal region of EMCV 2C is absolutely critical for EMCV 2C-mediated NDP52 degradation and autophagy induction.

**Fig 5.**
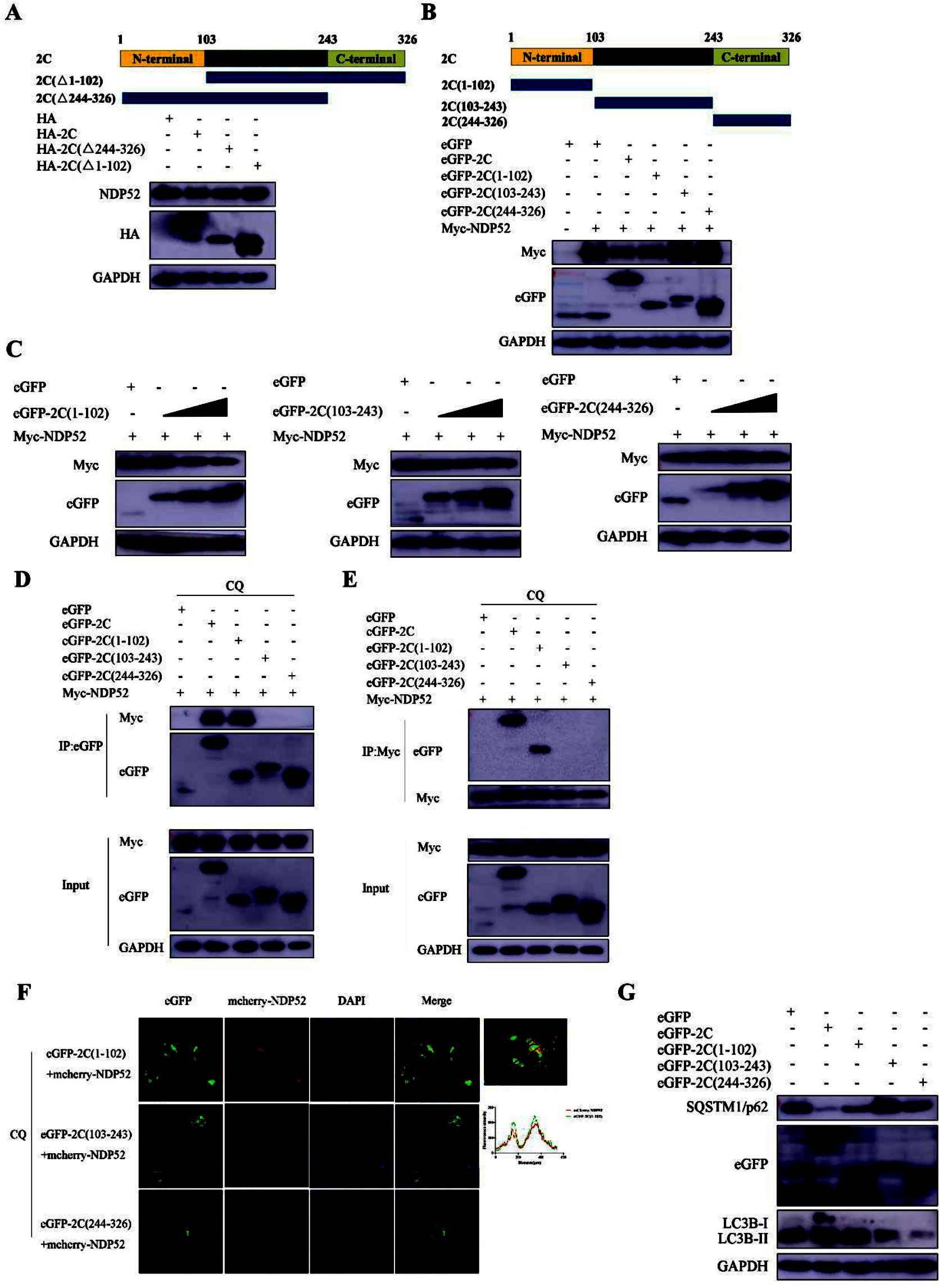
EMCV 2C interacts with and degrades NDP52 via the N-terminal region. (A) Schematic diagram of the structural domains and truncated 2C mutants. N-terminal: membrane-bound structural domain, ATPase domain: Activation of ATPase activity, C-terminal: deconjugating enzyme domain. A549 cells were transfected with 1.0 µg of HA or HA-2C, HA-2C (△1-102), HA-2C (△244-326) plasmids for 36 h. Samples were analyzed by western blotting with antibodies against NDP52, HA, and GAPDH. (B) A549 cells were cotransfected with eGFP-2C and its truncation constructs with Myc-NDP52 plasmids for 36 h. Samples were analyzed by western blotting with antibodies against GFP, Myc, and GAPDH. (C)A549 cells were cotransfected with 1.0 μg eGFP vector and eGFP-2C(1-102)/(103-243)/(244-326) (0.5 μg, 1.0 μg, 2.0 μg) plasmids with Myc-NDP52 for 36 h. Samples were analyzed by western blotting with antibodies against GFP, Myc, and GAPDH. (D, E)A549 cells were cotransfected with1.0 μg eGFP-2C and 1.0 μg its truncation constructs with 1.0 μg Myc-NDP52 plasmids for 24 h followed by CQ (50 µM). Samples were analyzed for Co-IP assay. (F)A549 cells were cotransfected with 1.0 μg mCherry-NDP52 and eGFP-2C(1-102) plasmids for 24 h. Samples were stained with DAPI (blue), then examined by confocal microscopy. Scale bar: 20 μm. (G) A549 cells were cotransfected with 1.0 μg eGFP vector and eGFP-2C(1-102)/(103-243)/(244-326) plasmids for 24 h. Samples were stained with DAPI (blue), then examined by confocal microscopy. Scale bar: 20 μm. (H) A549 cells were cotransfected with 1.0 μg eGFP vector and eGFP-2C(1-102)/(103-243)/(244-326) plasmids for 36 h. Western blotting was used to detect the protein expression of SQSTM1/p62, GFP, LC3B. GAPDH was used as a loading control.

### NDP52 expression dampens EMCV infection post-entry

NDP52-mediated selective autophagy is known to play an important role in regulating host antiviral responses (29). Indeed, EMCV viral titers were significantly lower in cells overexpressing NDP52 compared to control cells (Fig. 6A) correlating with a time-dependent decrease in EMCV VP1 expression and viral replication (Fig. 6B and C), an effect that can be reversed when NDP52 is knocked down (Fig. 6D and E). Moreover, we can demonstrate the antiviral effect in another cell line, HEK293 cells (Fig. 6F and G). These data highlight the clear antiviral nature of NDP52. Next, we wanted to know whether NDP52 functions in the early stages of infection. We found that NDP52 had little effect during viral adsorption phases, but inhibited EMCV during the viral entry stage (Fig. 6H and K), demonstrating that NDP52 functions during the intracellular viral invasion stage of infection.

**Fig 6.**
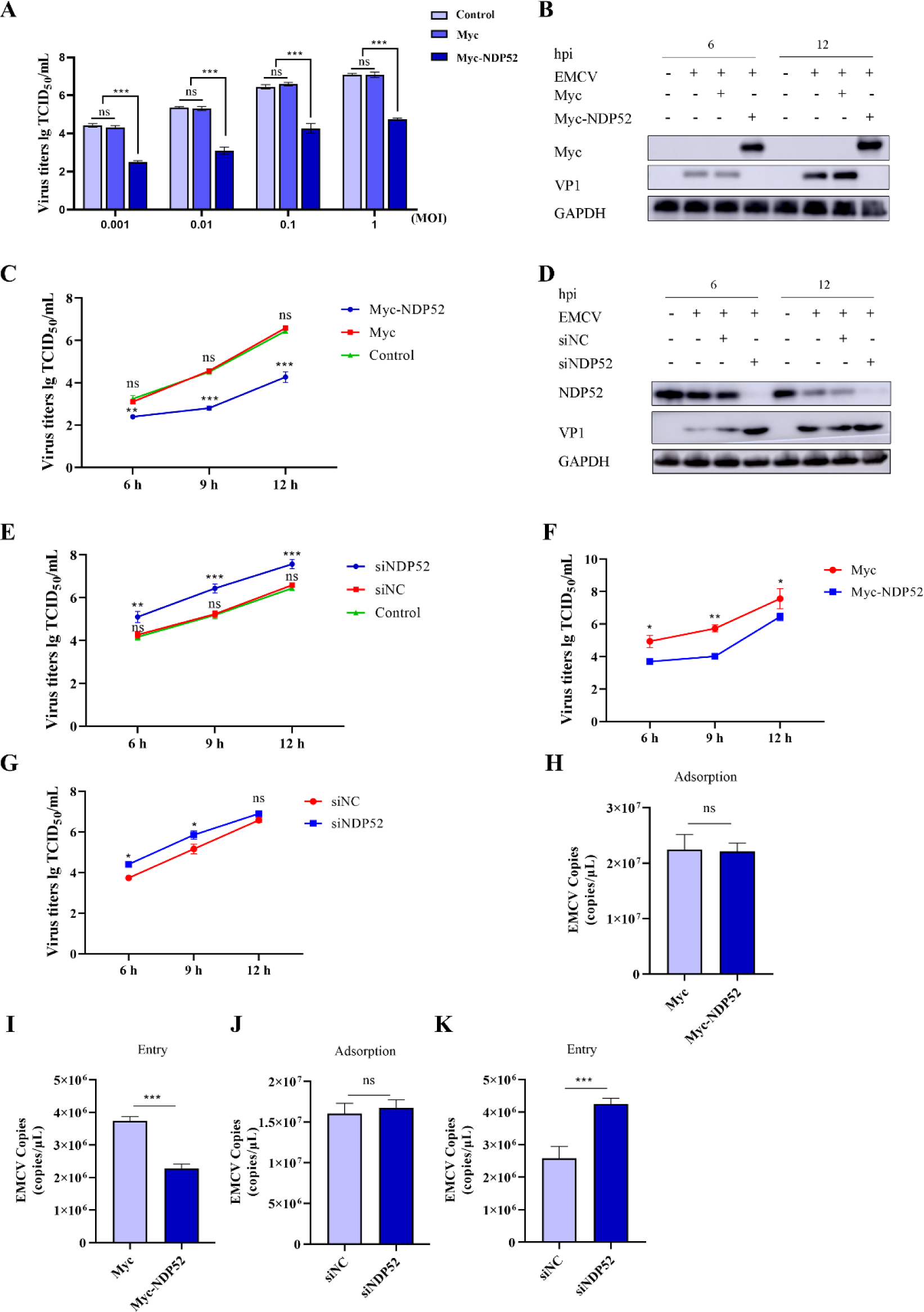
NDP52 expression dampens EMCV infection post-entry. (A) A549 cells were untransfected and transfected with 1.0 μg of EV, Myc-NDP52 for 36 h followed by EMCV (MOI =0.001/0.01/0.1/1) infection for 12 h. Samples were subjected to TCID_50_ assay. (B, C)A549 cells were untransfected and transfected with 1.0 μg of EV, Myc-NDP52 for 36 h followed by EMCV (MOI =1) infection for 6 h, 9 h and 12 h.Samples were subjected to western blot analysis and TCID_50_ assay, respectively. (D,E)A549 cells were transfected with 20mM of siNC or siRNA(siNDP52) for 36 h followed by EMCV (MOI = 1)infection for 6 h, 9 h and 12 h.Samples were subjected to western blot analysis and TCID_50_ assay, respectively. (F,G)HEK293 cells were transfected with 1.0 μg of EV, Myc-NDP52, or 20mM of siNC or siNDP52 for 36 h followed by EMCV (MOI = 1) infection for 6 h, 9 h and 12 h. Samples were subjected to TCID_50_ assay. (H,J) A549 cells were untransfected and transfected with 1.0 μg EV, Myc-NDP52 or 12 μl siNC, siNDP52 for 36 h followed by EMCV (MOI =3) infection for 1 h at 4℃. Then, the cells were washed 6 times with PBS. Virus RNA was extracted, and then the EMCV copies were detected by qRT-PCR. Statistical data from three independent infection experiments is shown.(I,K)A549 cells were untransfected and transfected with 1.0 μg EV, Myc-NDP52 or 12 μl siNC, siNDP52 for 36 h followed by EMCV (MOI =1) infection for 1 h at 4℃. Then, the cells were washed 6 times with PBS and continued to be added to the culture medium at 5% CO_2_ for 0.5 h and 1 h at 37°C. Virus RNA was extracted, and then the EMCV copies were detected by qRT-PCR. Statistical data from three independent infection experiments is shown. The data were represented as the mean ± SD of three independent experiments. The measurements were performed in technical duplicate. Statistical significance was denoted as \*\**P* < 0.01, and \*\*\**P* < 0.001.

### NDP52 targets EMCV VP1 and VP2 protein degradation via the autophagic lysosomal pathway

Autophagy receptors are known exert its antiviral functions by direct degradation of structural viral proteins (25). We wanted to explore the mechanism by which NDP52 degrades viral proteins encoded by EMCV, particularly its structural proteins VP1, 2 and 3. The exogenous co-expression of NDP52 with EMCV VP1, VP2 and VP3 resulted in the dose-dependent degradation of VP1 and VP2, not VP3, by NDP52 (Fig. 7A) underlining the selective nature of the autophagic degradation of EMCV proteins by NDP52. Moreover, we showed that NDP52 directly interacts with VP1 and VP2 but not VP3 (Fig. 7B-D). We further confirmed that NDP52 exerts its degradative function through the autophagic lysosomal pathway where VP1 and VP2 protein degradation can be blocked in the presence of an autophagy inhibitor (Fig. 7E and F). Importantly, exogenous introduction of NDP52 into cells promoted the autophagic conversion of LC3B-I to LC3B-II in a dose dependent manner (Fig. 7G) and that it co-localizes with LC3B structures in the cytosol (Fig. 7H). Overall, we show that NDP52 targets the degradation of EMCV VP1 and VP2 through the autophagic lysosomal pathway to inhibit EMCV proliferation.

**Fig 7.**
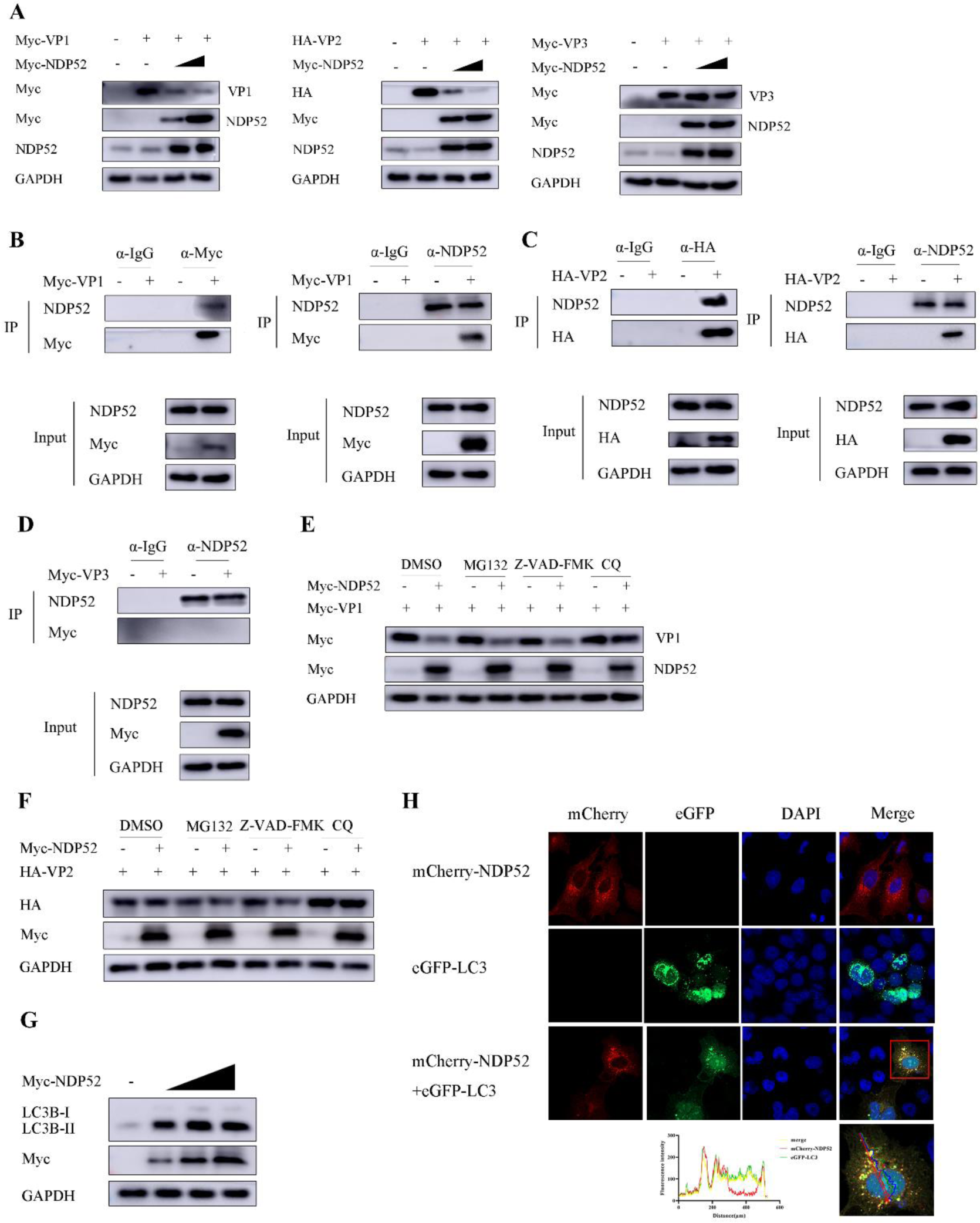
NDP52 targets EMCV VP1 and VP2 protein degradation via the autophagic lysosomal pathway. (A) A549 cells were transfected with 1.0 µg of EV or Myc-VP1, HA-VP2, Myc-VP3 for 36 h. Western blotting was used to detect the protein expression of Myc-NDP52, Myc-VP1, HA-VP2, Myc-VP3 proteins. GAPDH was used as a loading control. (B,C,D) A549 cells were transfected with 1.0 µg of EV or Myc-VP1, HA-VP2, Myc-VP3 for 36 h. Samples were analyzed for Co-IP assay. (E,F)A549 cells were co-transfected with 1.0 µg plasmid of EV and Myc-NDP52 or co-transfected with Myc-VP1/HA-VP2 and Myc-NDP52 plasmids for 24 h, and then treated for 8 h with DMSO, MG132 (10 µM), CQ (50 µM), or Z-VAD-FMK (20 µM), respectively. Western blotting was used to detect the expression of Myc-NDP52, Myc-VP1, HA-VP2 proteins. GAPDH was used as a loading control. (G) A549 cells were transfected with 1.0 µg of EV or Myc-NDP52 (0.5 µg, 1.0 µg, 2.0 µg) plasmids for 36 h. Western blotting was used to detect the expression of LC3B, Myc-NDP52 proteins. GAPDH was used as a loading control. (H) A549 cells were cotransfected with 1.0 μg mCherry-NDP52, eGFP-LC3 plasmids for 24 h. Samples were stained with DAPI (blue), then examined by confocal microscopy. Scale bar: 20 μm.

### 2C protein degrades NDP52 and relieves NDP52-mediated VP1 and VP2 degradation

Considering our observation that NDP52 inhibits EMCV infection early during the infection stages (Fig. 6), we wanted to explore whether EMCV 2C, which can directly degrade NDP52 (Fig. 3), is also involved in the early stages of EMCV infection. Indeed when we overexpressed 2C we observed enhanced adsorption and entry of EMCV (Fig. 8A and B) and this effect reversed when endogenous NDP52 was knocked down (Fig. 8C, D). From this, it is possible that EMCV 2C could be antagonizing NDP52-mediated degradation of VP1 and VP2. When EMCV 2C was overexpressed in a dose-dependent manner, we observed NDP52 degradation coinciding with the reversion of VP1 and VP2 levels (Fig. 8E and F), an effect that is pronounced in the presence of an EMCV infection (Fig. 8G). Our results indicate that 2C can degrade NDP52 whereby NDP52 loses its capacity to degrade VP1 and VP2.

**Fig 8.**
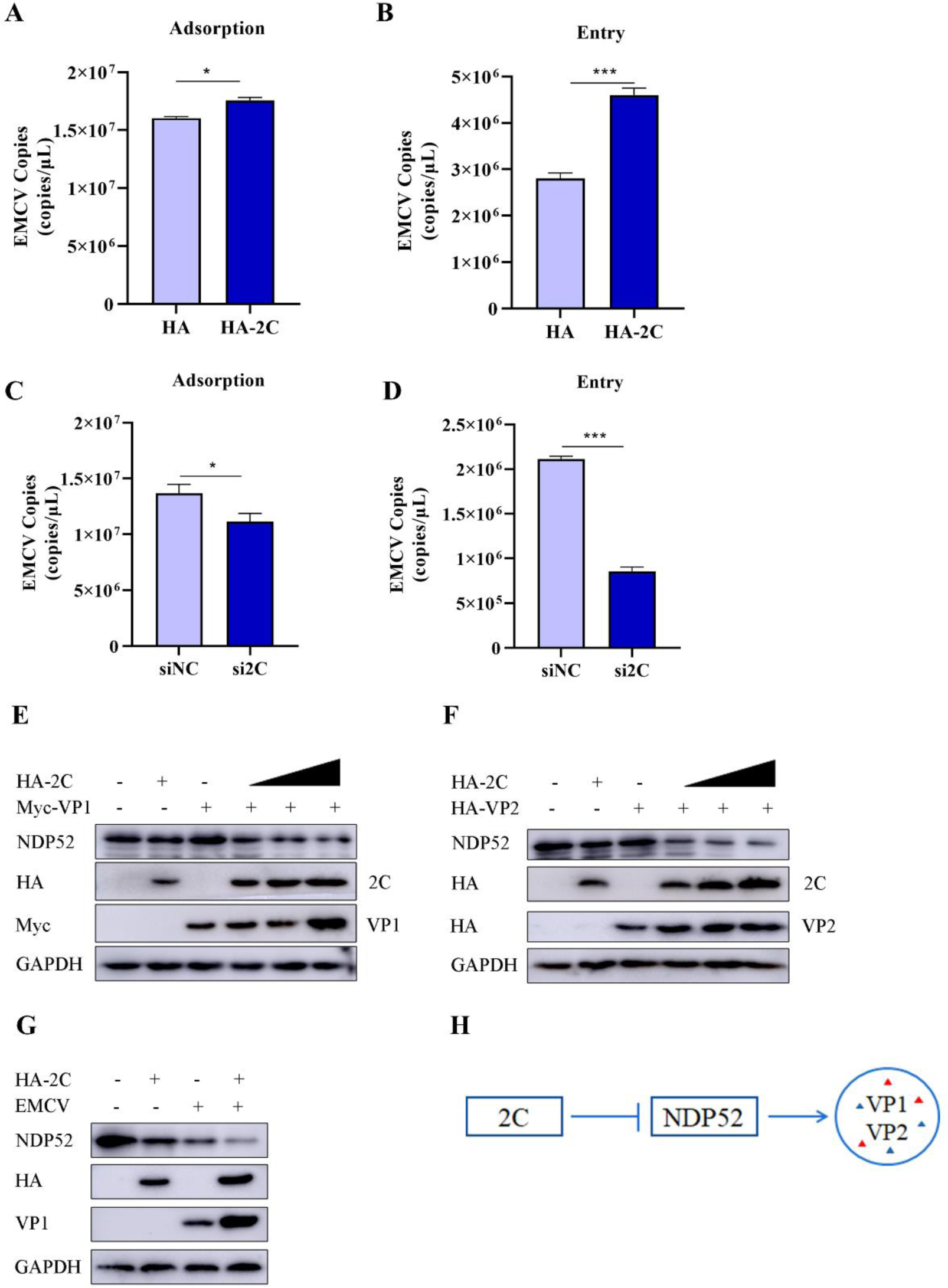
2C protein degrades NDP52 and relieves NDP52-mediated VP1 and VP2 degradation. (A, B) A549 cells were untransfected and transfected with 1.0 μg HA, HA-2C plasmids or 12 μl siNC, si2C RNA for 36 h followed by EMCV (MOI =3) infection for 1 h at 4℃. Then, the cells were washed 6 times with PBS. Virus RNA was extracted, and then the EMCV copies were detected by qRT-PCR. Statistical data from three independent infection experiments is shown. (C, D) A549 cells were untransfected and transfected with 1.0 μg HA, HA-2C plasmids or 12 μl siNC, si2C RNA for 36 h followed by EMCV (MOI =1) infection for 1 h at 4℃. Then, the cells were washed 6 times with PBS and continued to be added to the culture medium at 5% CO_2_ for 0.5 h and 1 h at 37°C. Virus RNA was extracted, and then the EMCV copies were detected by qRT-PCR. Statistical data from three independent infection experiments is shown.The data were represented as the mean ± SD of three independent experiments. The measurements were performed in technical duplicate. Statistical significance was denoted as \*\**P* < 0.01, and \*\*\**P* < 0.001. (E, F) A549 cells were transfected with 1.0 µg plasmid of EV and cotransfected with HA-2C (different dose) and Myc-VP1/HA-VP2 for 36 h. (G) A549 cells were transfected with 1.0 µg plasmid of HA and HA-2C for 36 h, then infected EMCV (MOI=1) for 8 h. Western blotting was used to detect the expression of NDP52 protein and VP1. GAPDH was used as a loading control.

## Discussion

Autophagy receptors can be classified as non-selective and selective autophagy based on their involvement in specific recognition binding to substrates (30). To date, most of the mechanistic studies on selective autophagy suggest that autophagy receptors are involved in the recognition and binding of relevant target proteins, including viral proteins in the cytoplasm, which then interact with LC3 to form a double-membrane structure to form autophagosomes destined for lysosomal degradation. Optineurin (OPTN), a key regulator of autophagy, was shown to target SVV VP1 for degradation to inhibit viral infectivity (25). The N protein from several coronaviruses can undergo ubiquitination and direct lysosomal degradation by NDP52 to inhibit coronavirus replication (31). Recent studies also show that NDP52 can regulate the process of viral endocytosis. NDP52 was shown to inhibit viral proliferation by degrading the M protein of hepatitis B (HBV) in an autophagy-independent manner, but through Rab9-mediated lysosomal pathway targeting at the viral internalization phase (22). In our present study, we also found that NDP52 inhibited EMCV internalization, followed by EMCV VP1 and VP2 targeting for autophagic degradation, to restrict EMCV replication.

NDP52 is made up of several domains, in particular an atypical LC3 interaction region (LIR) motif that specifically interacts with LC3, which is critical for NDP52 attachment to autophagosomes, a cargo-binding region (PPI) consisting of a Galectin 8-interacting region (GIR), which is important for recognizing substrate proteins and a zinc finger structural domain (ZF) that recognizes a number of ubiquitin proteins (32, 33). It is unsurprising that viruses have evolved strategies to interfere with the host autophagy machinery by antagonizing autophagy receptors such as NDP52, or by holding autophagy receptors hostage and in turn degrading host immune proteins to accomplish immune escape. For example, coxsackievirus B3 (CVB3) can evade heterophilic autophagy by cleaving and degrading NDP52 post-infection infection and holds NDP52 hostage to relieve its interaction with CVB3 VP1 protein. CVB3 then inhibits type I interferon (IFN) responses by the promoting autophagic degradation of mitochondrial anti-viral signaling proteins (MAVS) to promote its own proliferation (29). Similarly, influenza A NP protein has been shown to interact with MAVS and Toll-interacting protein (TOLLIP), another known autophagy regulator, whereby it holds TOLLIP hostage to degrade MAVS to eventuate in the blockade of antiviral signaling to promote viral replication (34).

The interplay between EMCV and the autophagy machinery was previously reported whereby EMCV 2C and 3D proteins induces autophagy and promotes viral replication by initiating the unfolded protein response (UPR) pathway in response to endoplasmic reticulum (ER) stress (8, 35). Our study also describes the existence of a reciprocal ‘check and balance’ between EMCV and the autophagic machinery, where EMCV induces autophagy through its encoded 2C, which degrades the autophagy receptor NDP52 to escape the autophagic antiviral response. Moreover, we describe the key involvement of the 2C N-terminal region to not only promote autophagy, but to participate in the direct interaction with NDP52 to drive its degradation. Importantly, we showed that this NDP52 degradation effect occurs exclusively with 2C protein from EMCV. Rab proteins are small GTPases belonging to the Ras-like GTPase superfamily that regulate vesicular transport. Many Rab proteins have been shown to be involved in various stages of autophagy (18). Rab7 plays a key role in autophagosome maturation, whereas Rab9 is a critical protein in the non-classical autophagy pathway (21). In our gene silencing experiments, we demonstrated the crucial role of Rab7 and Rab9 in EMCV-induced autophagy-lysosomal degradation of NDP52.

In conclusion, we describe a novel role for NDP52 in anti-EMCV defense. We show that NDP52 can inhibit EMCV replication by modulating early EMCV endocytosis and by controlling autophagic degradation of EMCV structural proteins VP1 and VP2. EMCV then confers it evasive strategy by exerting its autophagy-inducing function through its own encoded 2C protein by direct interaction with NDP52. 2C then leads NDP52 to lysosomal-mediated degradation, a process relying on the involvement of late endosomal Rab7 and Rab9 proteins, presumably playing an important role in transporting NDP52 to the lysosome. These findings provide further insight into the relationship between EMCV and autophagy and provide additional evidence for the role of autophagy receptors in antiviral defense. Our study also provides new evidence of viral escape strategies against the autophagy machinery and insights into potentially new antiviral therapeutic targets.

## Materials and methods

### Cells and Viruses

Human non-small cell lung cancer cells (A549) cells were cultured in F-12 (Cellmax, Lanzhou, China) supplemented with 15% new bovine serum (NBS) (Lanzhou Minhai, Lanzhou, China). Human embryonic kidney (HEK293) cells and Baby hamster syrian kidney (BHK-21) cells were cultured in DMEM (Cellmax, Lanzhou, China) supplemented with 10% new bovine serum (NBS) (Lanzhou Minhai, Lanzhou, China), They were obtained from the American Type Culture Collection (ATCC) and maintained at a 37°C incubator containing 5% CO_2_. Encephalomyocarditis virus (EMCV) strain PV21 (GenBank No: X74312) was provided by Biomedical Research Center of Northwest Minzu University.

### Plasmids, Antibodies and Reagents

Expression plasmids mCherry-NDP52, eGFP-2C(103-243), and eGFP-2C(244-326) were synthesized by Public Protein/Plasmid Library (Nanjing, Jiangsu, China). Myc-NDP52, HA-2C(△1-102), HA-2C(△244-326), eGFP-2C, eGFP-2C(1-102), the HA-tagged 2C of enterovirus 71 (HA-2C EV-71), the HA-tagged 2C of foot-and-mouth disease virus (HA-2C FMDV), and the HA-tagged 2C of Seneca Valley virus (HA-2C SVA) plasmids were synthesized by Genscript (Nanjing, Jiangsu, China). Myc-VP1, HA-VP2, Myc-VP3, HA-2A, HA-2C, HA-3A, Myc-3C, and eGFP-LC3 plasmids were constructed in our laboratory.

Anti-NDP52 rabbit polyclonal antibody (pAb), anti-HA Tag mouse monoclonal antibody (mAb), anti-HA rabbit pAb, anti-Myc mouse mAb, anti-LC3B rabbit pAb, anti-GFP mouse mAb, and anti-GAPDH mouse mAb were purchased from Proteintech (Wuhan, Hubei, China). Rab7 Rabbit mAb and Rab9A Rabbit mAb were purchased from Zenbio (Chengdu, China). Anti-EMCV VP1 mouse pAb was provided by Gene Create (Wuhan, Hubei, China).

Lipofectamine 2000 and Lipofectamine 3000 were purchased from Invitrogen (Invitrogen, Waltham, MA, USA). MG132, NH4Cl and Caspase inhibitor Z-VAD-FMK were purchased from Beyotime (Shanghai, China). Chloroquine (CQ) was purchased from InvivoGen (San Diego, CA, USA). 3-Methyladenine and Bafilomycin A1 were purchased from TOPSCIENCE (Shenzhen, China). Hoechst 33342 and Lyso-Tracker Green were purchased from Solarbio (Beijing, China). Cycloheximide (CHX) was purchased from MedChemExpress (Monmouth Junction, NJ, USA).

### Cell transfection and Western blotting

A549 cells, HEK293 cells and BHK-21 cells in 6 well plates or 10 cm dishes were transfected with indicated expression plasmids using Lipofectamine 2000 (Invitrogen, Waltham, MA, USA) for 36h. Then, re-infected with EMCV for different times or added inhibitor treatment for 12h. Cells were collected and lysed with RIPA buffer (Beyotime, Shanghai, China) containing protease inhibitor and EDTA (Beyotime, Shanghai, China). Equal amounts of cell extracts were denatured, assayed on 10% or 12% SDS-PAGE gels, and then the proteins were transferred to PVDF membranes (Millipore, MA, USA) using a semi-dry transmembrane apparatus. The membranes were blocked using 2.5% skimmed milk plus TBST (Solarbio, Beijing, China) for 2h at room temperature, and then incubated with specific primary antibody (1:1000) for 12 h at 4°C and secondary antibody (1:10,000) for 1h at room temperature. Finally, proteins on membranes were detected using Western lightning Plus-ECL kit (PerkinElmer, Waltham, MA, USA).

### Virus infectivity assays

For in vitro virus infection, treated or untreated HEK293 cells or A549 cells were washed three times with PBS (Solarbio, Beijing, China) and infected with EMCV strain PV21 (MOI = 0.1/1/2/3) for different times. These cells were then continued to be cultured in maintenance medium for a certain period of time. The viral suspension was collected after freeze-thawing for three times, and the titers of different viruses were measured in BHK-21 cells by TCID_50_ method (Reed-Muench method).

### RNA extraction and quantitative PCR (qPCR)

Total intracellular RNA was extracted with TRIZOL reagent (Takara, Beijing, China). The extracted RNA was synthesized into cDNA using the Evo M-MLV Reverse Transcription Premix Kit (Accurate Biology, Hunan, China). The mRNA expression levels of *NDP52,* and *2C* were detected using TransStart Top Green qPCR SuperMix (+Dye II) (Transgen, Beijing, China). The relative abundances of target genes were normalized to that of GAPDH by the 2^−ΔΔCt^ method. The qPCR primers are shown in Table 1.

**Table 1.**
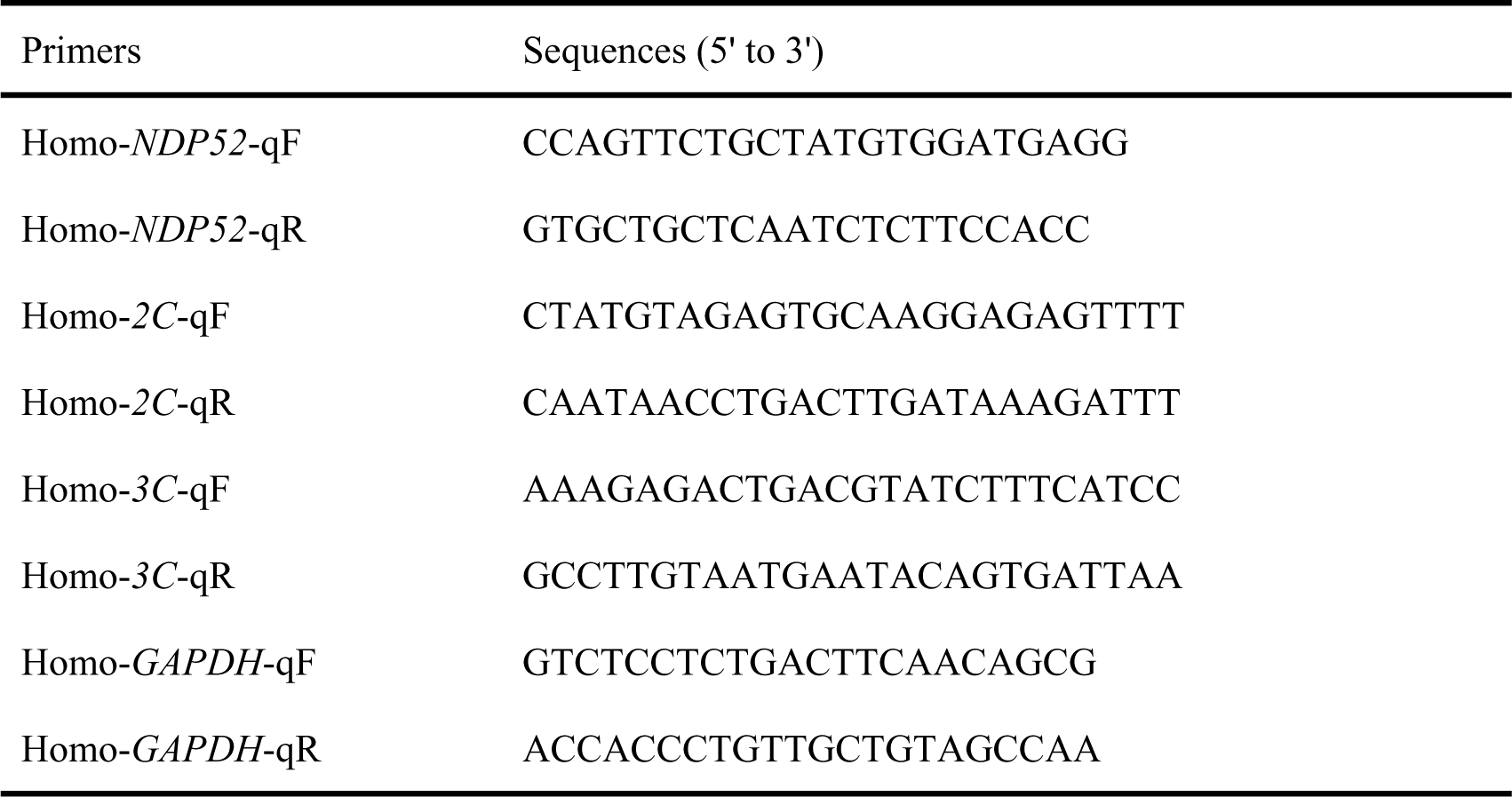
The sequences of qPCR primers used in this study.

### siRNA interference assay

When the density of A549 cells reached 30-40%, siRNA oligos was transfected into the cells with Lipofectamine 2000, and siNC was used as the negative control. siRNA was synthesized by RiboBio (Guangzhou, China). The siRNA sequences are shown in Table 2.

**Table 2.**
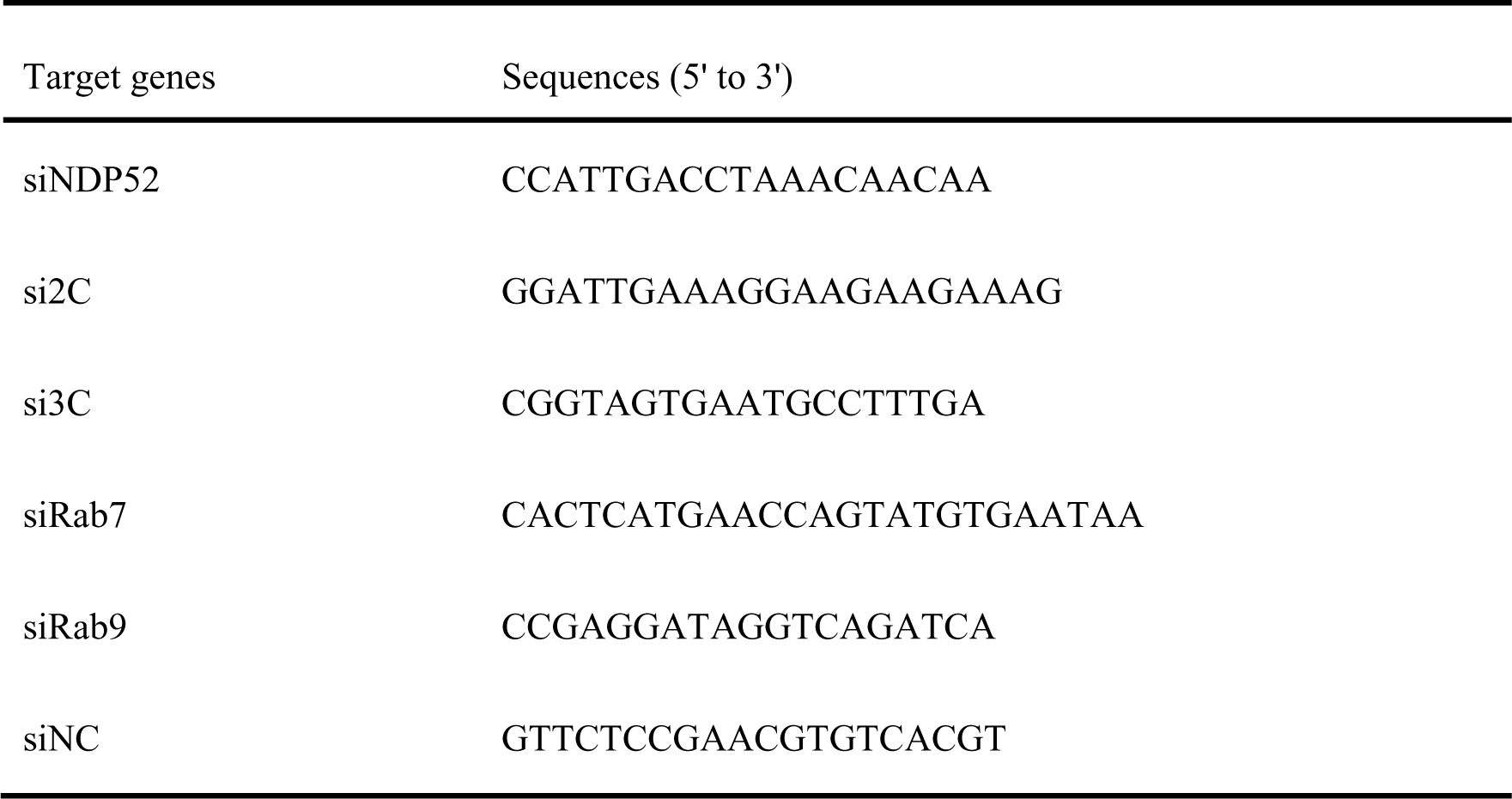
The siRNA sequences used in this study.

### Co-immunoprecipitation (Co-IP)

The transfected cells were collected and lysed using NP-40 buffer containing protease inhibitors (Beyotime, Shanghai, China). A portion of the cell lysate was used for Input following the protein immunoblotting method described above. Another portion was used as IP to adsorb the proteins in the cell lysate using protein G (Beyotime, Shanghai, China), followed by immunoprecipitation using the indicated antibodies. Immunoprecipitated samples was gathered and utilized for Western blotting.

### Inhibitor treatment assay

A549 cells in 6-well plates were transfected with pCMV-Myc (EV), Myc-NDP52 and other recombinant plasmids using Lipofectamine 2000. At 24 h post-transfection, the cells were treated with pan-caspase inhibitor Z-VAD-FMK (Beyotime, Shanghai, China), proteasome inhibitor MG132 (Beyotime, Shanghai, China), endosomal acidification and autophagy inhibitor CQ (InvivoGen, San Diego, CA, USA), or DMSO for 8 h, respectively. Then, the cells were collected and lysed with RIPA lysis buffer supplemented with protease and phosphatase inhibitors. Equal amounts of cell extracts were denatured and analyzed using 10% SDS-PAGE gels, and then transferred to methanol-activated PVDF (Millipore, MA, USA) membranes. Subsequently, these membranes were blocked 2.5% skim milk before specific primary and secondary antibodies were added, respectively. Finally, proteins on membranes were detected using ECL Substrate.

### Indirect immunofluorescence (IFA)

Cells cultured on Nunc glass-bottom petri dishes (ThermoFisher Scientific, Waltham, MA, USA) were transfected with various plasmids or infected with EMCV. Cells were fixed with 4% paraformaldehyde for 10 min and blocked with 1% PBST bovine serum albumin for 6-8 h at 4°C. Cells were then incubated with a 1:250 dilution of appropriate primary antibody overnight at 4°C, followed by incubation with a 1:500 dilution of Cy3-conjugated goat anti-mouse IgG and/or CoraLite488-conjugated goat anti-rabbit IgG (Jackson ImmunoResearch Laboratories, West Grove, PA, USA) for 1 h at room temperature in the dark. Cells were then stained with 4’,6-diamino-2-phenylindole (DAPI) for 10 min at room temperature to reveal the nuclei. Fluorescence was observed by ZEISS LSM900 laser confocal microscopy.

### Lysosome colocalization

A549 cells were transfected with the expression plasmids and after 12 h of cell culture,then infection EMCV(MOI=3) for 3 h and 6 h, and stained with Lyso-Tracker Green (Solarbio, Beijing, China) for 1h.And the cells were washed 3 times with PBS, Hoechst 33342 (Solarbio, Beijing, China) was then added to stain the nuclei for 10 min to reveal the nuclei. The fluorescence was observed by ZEISS LSM900 laser confocal microscope.

### Statistical Analysis

Data are expressed as the mean ± SD (standard deviation). Statistical significance was determined by using Student’s two-tailed non-parametric t-test or ANOVA (analysis of variance) with GraphPad Prism software (version 9.0, USA). Differences between groups were considered significant when the *P* value was < 0.05 (*), < 0.01 (**), and < 0.001 (***).

## CRediT authorship contribution statement

Rongqian Mo, Conceptualization, Data curation, Formal analysis, Investigation, Methodology, Software, Visualization, Writing-original draft; Rongrong Cheng, Data curation, Investigation, Methodology, Software; Pingan Dong, Investigation, Visualization; Tingting Ma, Investigation, Visualization; Yaxin Zhang, Investigation, Validation; Jingying Xie, Resources; Shasha Li, Methodology; Huixia Li, Investigation; Adi Idris, Writing-review & editing; Xiangrong Li, Conceptualization, Data curation, Funding acquisition, Methodology, Project administration, Resources, Supervision, Validation, Writing-review & editing; Ruofei Feng, Conceptualization, Formal analysis, Funding acquisition, Project administration, Supervision, Writing-review & editing.

## Data availability

The data that support the findings of this study are available from the corresponding authors upon request.

